# Local wavefront alignment with corrected free-end penalty

**DOI:** 10.64898/2026.11.23.768503

**Authors:** Jingwei Li, Haitao Zhu, Liling Jia, Qiang Wu

**Affiliations:** Center for Comparative Biomedicine, Key Laboratory of Systems Biomedicine (MOE), Institute of Systems Biomedicine, Shanghai Jiao Tong university, Dongchuan Road, Shanghai, 200240, Shanghai, China; Shanghai Key Laboratory of Gene Editing and Cell-based Immunotherapy for Hematological Diseases, State Key Laboratory of Medical Genomics, Ruijin Hospital, Shanghai Jiao Tong University School of Medicine, South Chongqing Road, Shanghai, 200025, Shanghai, China

**Keywords:** exact alignment, global alignment, local alignment, pairwise alignment, wavefront alignment

## Abstract

**Background:** Pairwise alignment of nucleotide and protein sequences is a fundamental problem in bioinformatics. Wavefront alignment (WFA) is a groundbreaking algorithmic advance that reduces the time complexity of exact (semi-)global pairwise alignment with affine gap penalties from ***O*(*N***^***2***^) to ***O*(*sN*)**. Despite its success, WFA has yet to be extended to the local sequence alignment scope.

**Results:** In this work, we propose a local wavefront alignment (lWFA) which generalizes WFA to the local alignment scope and preserves both the ***O*(*sN*)** time and the ***O*(*s***^***2***^) space complexity. We also propose the bidirectional lWFA (lBiWFA), which reduces the space complexity of lWFA from ***O*(*s***^***2***^) to ***O*(*s*)** while maintaining the time complexity as ***O*(*sN*)**. Benchmarks show that lWFA and lBiWFA outperform current exact local alignment libraries like SeqAn and Parasail in aligning long sequences with error rate less than **5%**.

**Conclusions:** We suggest to use lBiWFA over lWFA because it has negligible memory footprint for aligning reads up to 20kbp, and is faster than lWFA except for short reads with very low error rates. According to the real-data benchmarks, lBiWFA are faster than SeqAn and Parasail for aligning NGS short reads with 0.1%∼1% error rate and ONT long reads with 3%∼5% error rate. However, for reads sequenced by legacy long-read platforms with 10%∼15% error rate, SeqAn and Parasail are faster. Nevertheless, lBiWFA has a competitive speed for reads with error rate up to 10%. One may choose lBiWFA over SeqAn and Parasail if the memory usage is the major concern.

## 1 Background

Pairwise alignment is a fundamental building block for biological sequence analyses. It underlies reads mapping [1], *de novo* genome assembly [2], variants calling [3], multiple sequence alignment [4], and many other applications. Given its ubiquitous importance, pairwise alignment has been continuously studied for decades and remains an active area of research. With the advent of high-throughput second- and third-generation sequencing technologies, the volume of sequencing data has increased dramatically and sequences to be aligned have grown longer. For example, high-throughput gene editing and 3D genomics has generated voluminous sequencing datasets that need to be aligned rapidly with references [5, 6]. This underscores the need for faster algorithms capable of keeping pace with growing data volumes and adapting to the longer reads produced by modern sequencing platforms.

Pairwise alignment algorithms fall into two categories: heuristic and exact. Heuristic methods, such as BLAST [7] and Paraband [8], may produce suboptimal alignments. In contrast, exact methods like the Needleman–Wunsch (NW) [9] and the Smith–Waterman (SW) [10] algorithms always guarantee the optimal alignment. These classical exact approaches employ dynamic programming (DP) to construct a score table of size *O*(*N*^2^), which can then be traced back to obtain the optimal alignment. Although various acceleration techniques and heuristics, including Single Instruction, Multiple Data (SIMD) [11], prefix-scan [12], and bit-vector methods [13], have been developed, the quadratic time complexity of classical DP-based exact methods remains a major bottleneck, limiting their applicability to long reads.

Wavefront alignment (WFA) is a groundbreaking method for pairwise alignment with affine gap penalties [14]. As an exact method, WFA always finds the optimal alignment. It adapts Myers’s *O*(*ND*) algorithm [15] to the widely-used affine gap penalties [16]. Instead of computing the scores of partial alignments, WFA fixes the score and attempts to extend the alignment as far as possible. WFA achieves an *O*(*sN* ) time complexity instead of *O*(*N*^2^), making it suitable for aligning sequences with moderate error rates (a common scenario in practice).

Since the introduction of WFA, many subsequent studies have continued to improve upon it. López-Villellas *et al*. optimize the space usage of the backtrack process in WFA [17]. Hardware-based accelerations, such as those using GPUs and FPGAs, have also been implemented for WFA [18, 19]. Eizenga and Paten improve the asymptotic space and time complexity of WFA from *O*(*s*^2^) and *O*(*sN* ) to *O*(*s*^3*/*2^) and *O*(*s*^2^ + *N* ), respectively [20]. Through a divide-and-conquer design, the bidirectional WFA algorithm (BiWFA) [21] reduces the space complexity of WFA from *O*(*s*^2^) to *O*(*s*) [22, 23].

In this work, we adapt the wavefront scoring system to the local alignment scope by introducing linear penalties to unaligned regions. We prove that our adapted wavefront scoring system is equivalent to the classical affine gap scoring system. Based on this scoring system, we propose the wavefront alignment for the local scope (lWFA) with *O*(*s*^2^) space complexity and *O*(*sN* ) time complexity, as well as its bidirectional version (lBiWFA) with *O*(*s*) space complexity and *O*(*sN* ) time complexity. Our benchmarks demonstrate that lBiWFA outperforms existing exact local alignment libraries such as Parasail and SeqAn in both time and space efficiency when aligning reads with moderate error rates.

## 2 Methods

We give strict theories in this section. For a non-strict yet intuitive walkthrough of our core ideas, readers may skip ahead to subsection 3.1.

### 2.1 Scoring system for local wavefront alignment

We use similar symbols as in previous works [14, 20, 21]. Denote the text and query by *t* and *q* with lengths *M* and *N*, respectively. For each alignment, define [20]:

- *L* the total length of matches;
- *X* the total length of mismatches;
- *O* the total number of gaps;
- *E* the total length of gaps.

Global WFA uses a scoring system [14] different from the classical one [16]. It penalizes

- mismatch by *x >* 0;
- gap opening by *o >* 0;
- gap extension by *e >* 0.

No score is awarded for matches. WFA searches an alignment minimizing the score

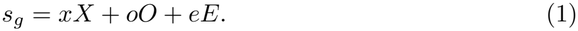

The WFA scoring system does not make sense in the local alignment scope because one always achieves the overall minimal (optimal) score of zero by simply not aligning at all.

The global WFA scoring system has only a superficial difference from the classical scoring system [20], defined as:

- *l*_*c*_ *>* 0 the match score;
- *x*_*c*_ *>* 0 the mismatch penalty;
- *o*_*c*_ *>* 0 the gap opening penalty;
- *e*_*c*_ *>* 0 the gap extension penalty.

They set the WFA scoring system as

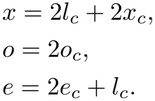

By the identity

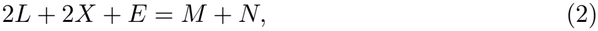

the authors show that the classical score

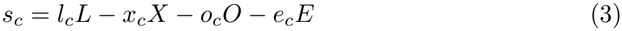

and the global WFA score *s*_*g*_ in Eq. (1) are related by

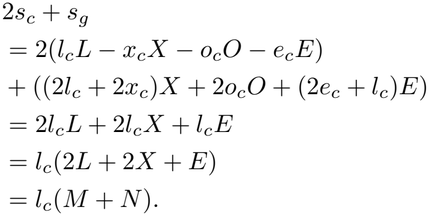

This implies that maximizing the classical score *s*_*c*_ is equivalent to minimizing the WFA score *s*_*g*_.

The key component to relate these two scoring systems in the global alignment scope is the identity in Eq. (2). For the local alignment scope, Eq. (2) is no longer valid because only a local region with undetermined size is aligned. To construct a similar identity for the local alignment scope, define *J* as the total size of unaligned regions at both ends in the local alignment. A local alignment can generate an unaligned region at the start by jumping in the DP table from cell (0, 0) to cell (*m*_0_, *n*_0_). Then it grows the aligned region from cell (*m*_0_, *n*_0_) to cell (*m*_1_, *n*_1_). Finally, an unaligned region at the end is generated by jumping from cell (*m*_1_, *n*_1_) to cell (*M, N* ). In this case,

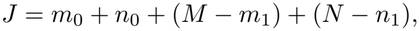

and we have the identity for the local scope

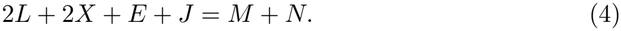

Define *j* as the penalty for each base in the unaligned regions at both ends. The lWFA minimizes the score

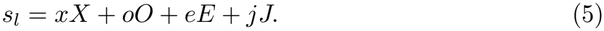

The classical score *s*_*c*_ of the local alignment scope is the same as that of the global one in Eq. (3) because the classical scoring system does not penalize unaligned regions. Let

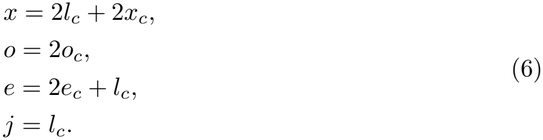

Then *s*_*c*_ and *s*_*l*_ are related by

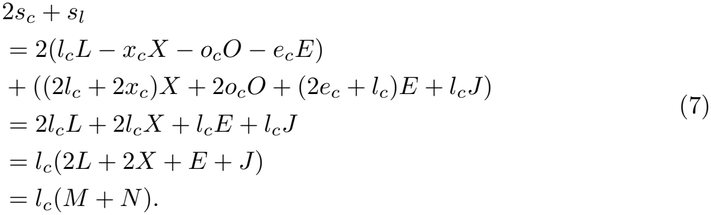

The lWFA scoring system in Eq. (6) has the following intuitive explanation.

- A mismatch not only has a penalty *x*_*c*_ in the classical scoring system, but also wastes a chance for the match. Thus, *x* = *l*_*c*_ + *x*_*c*_.
- The gap opening penalty is simply passed forward by *o* = *o*_*c*_.
- A gap extension not only has a penalty *e*_*c*_ in the classical scoring system, but also wastes half a chance for the match. It is half a waste because a match increases the offset by 2 (1 in each of the query and text dimensions in the DP table), while a gap extension only increases the offset by 1. Thus, *e* = *e*_*c*_ + *l*_*c*_*/*2.
- A jumping step wastes half a chance for the match. Thus, *j* = *l*_*c*_*/*2.
- To keep the scores as integers, double the whole scoring system by a factor of 2, which leads to Eq. (6).

The WFA implementation in WFA2-lib includes several semi-global alignment spans [14, 21]. WFA2-lib defaults to zero penalties for free ends in the semi-global alignment scope, as shown in Eq. (8).

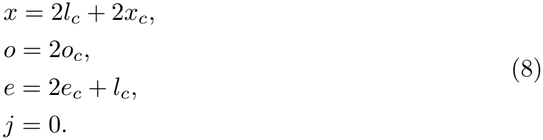

WFA2-lib minimizes the target score *s*_*g*_ in Eq. (1) for both the global and semi-global alignment scope. The WFA2-lib semi-global scoring system in Eq. (8) is different from the classical semi-global scoring system. Instead, it is equivalent to the modified version of the classical semi-global scoring system in Eq. (9) which awards the free ends by *l*_*c*_*/*2.

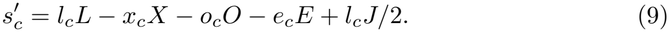

Actually,

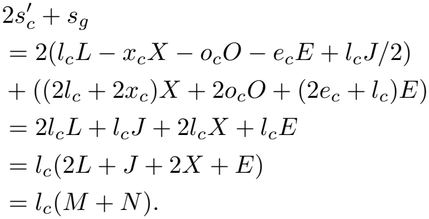

### 2.2 The local wavefront alignment

With the same symbols used in previous works [20, 21], we define the wavefront components for the local alignment scope:

- *ℐ*_*s,d*_ the maximal offset of the partial alignments ending at diagonal *d* with score *s* and a tailing insertion.
- *D*_*s,d*_ that with a tailing deletion.
- *X*_*s,d*_ that with a tailing insertion, deletion, mismatch or jumping.
- ℳ_*s,d*_ that with a tailing perfect matching extension from *X*_*s,d*_ along diagonal *d*.

The above definitions are different from the global alignment scope in that *X*_*s,d*_ may terminate with a jumping. Define *µ*(*m, n*) as the length of the longest matching prefix between *t*_*m*:*M*_ and *q*_*n*:*N*_ . The recurrence of the wavefront components is

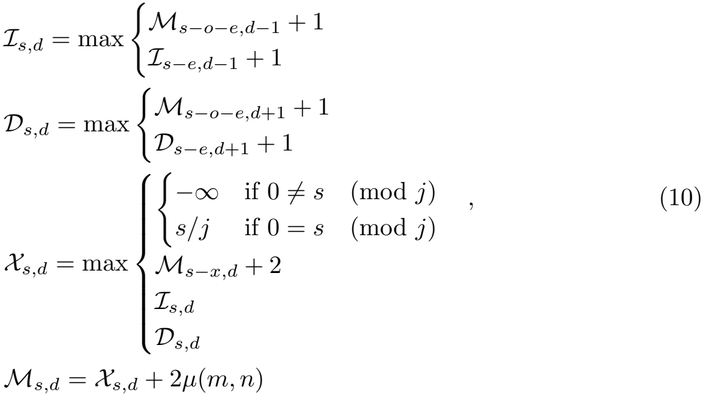

where (*m, n*) is the DP cell of *X*_*s,d*_ with

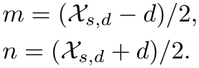

In the recurrence of *X*_*s,d*_, if 0 = *s* (mod *j*), then a jumping of size *f* = *s/j* has a score exactly equal to *s*, thereby forcing a lower bound *f* for *X*_*s,d*_ on all diagonals *d* satisfying *d* = *f* (mod 2). We prove the correctness of Eq. (10) by a modified version of LEMMA 2.1 in [14]. The main difference from the original proof is that we include the discussion of jumping (refer Appendix A.1 for the proof).

#### Lemma 1

*The following is valid in Eq. (10)*.

- *ℳ*_*s,d*_ *is the maximal offset of the partial alignments ending at diagonal d with score s*.
- ℐ_*s,d*_ *is that with a tailing insertion*.
- *D*_*s,d*_ *is that with a tailing deletion*.

The diagonal range of lWFA expands towards both sides as the wave score *s* increasing due to the insertion and deletion. For *C* ∈ {*D, ℐ, X*, ℳ}, define diag(*C*_*s*_) as the set of diagonals *d* such that *C*_*s,d*_ is involved in the recurrence in Eq. (10). Algorithm 1 gives a unified implementation of the WFA for both the global and local alignment scopes. Algorithm 1 introduces the following new symbols. The optimal score so far is stored in *s*_*t*_. Let 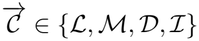 with

- 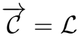 the alignment with a heading jumping;
- 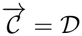 the alignment with a heading deletion;
- 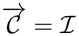the alignment with a heading insertion;
- 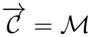 the alignment not with a heading jumping.

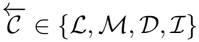 is similar to 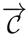 but restricts the alignment tailing instead of heading. By properly setting 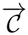 and 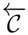, Algorithm 1 can adapt to various alignment scopes, including the global and local scopes. This is especially useful for the implementation of lBiWFA.

In Algorithm 1, we have used the convention that if *C*_*s,d*_ with *C* ∈ {*D*, ℐ, *X*, ℳ} does not exist when accessing, then let *C*_*s,d*_ = −∞. The correctness of Algorithm 1 is given in Proposition 1 (refer Appendix A.2 for the proof).

#### Proposition 1

*Algorithm 1 returns the optimal alignment*.

Algorithm 1 preserves the same time and space complexity as the global version (refer Appendix A.3 for the proof).

#### Proposition 2

*The time and space complexity of Algorithm 1 is O*(*s*_*o*_(*M* + *N* )) *and* 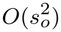, *respectively, where s*_*o*_ *is the optimal alignment score*.

### 2.3 The local bidirectional wavefront alignment

In [21], the authors propose BiWFA based on a divide-and-conquer method, which reduces the space complexity of global WFA from *O*(*s*^2^) to *O*(*s*). We propose a similar method lBiWFA for lWFA in Algorithm 2 with the following additional symbols.

#### Algorithm 1

The unified WFA for the global and local alignment scopes.

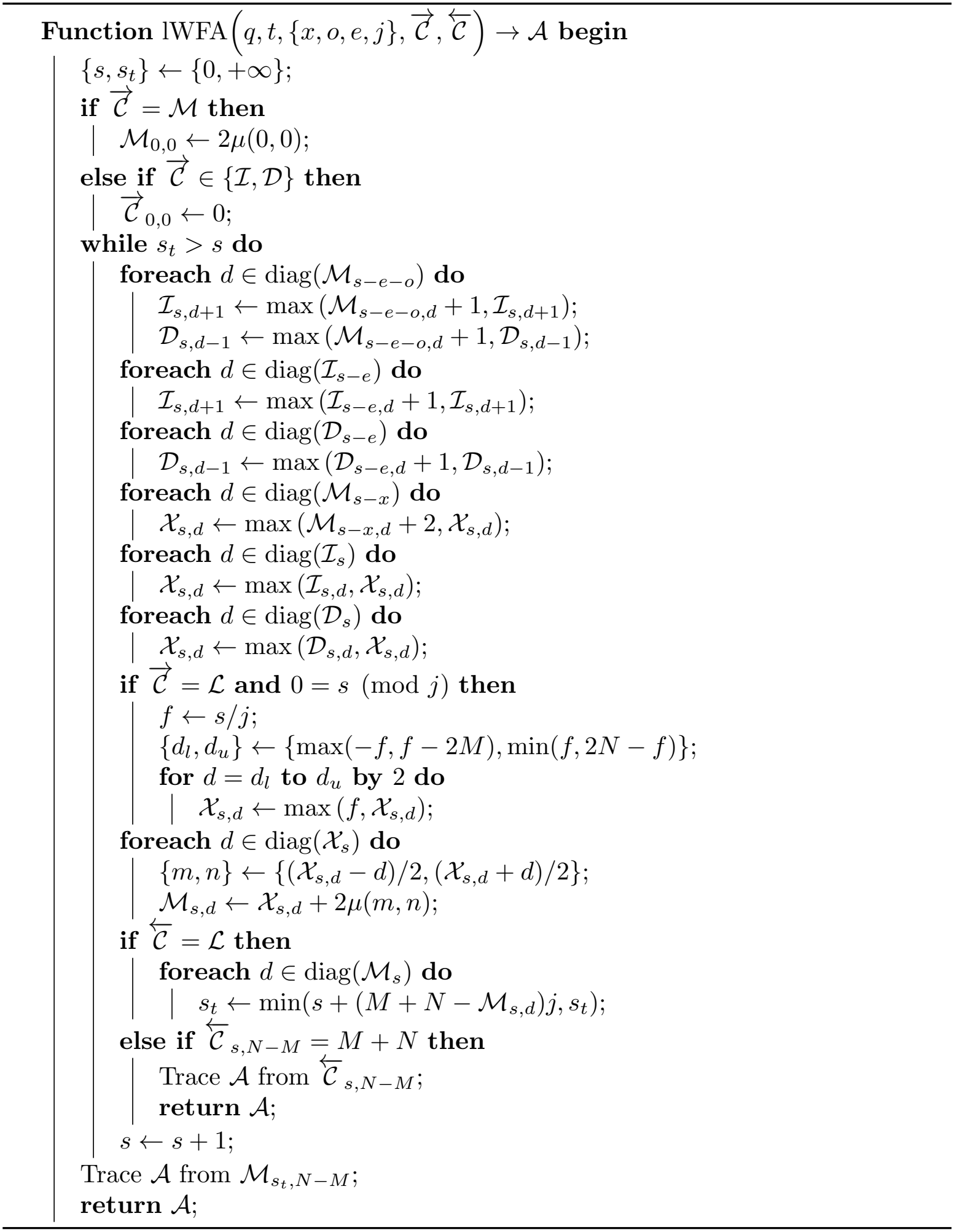

#### Algorithm 2

The unified BiWFA for the global and local alignment scopes.

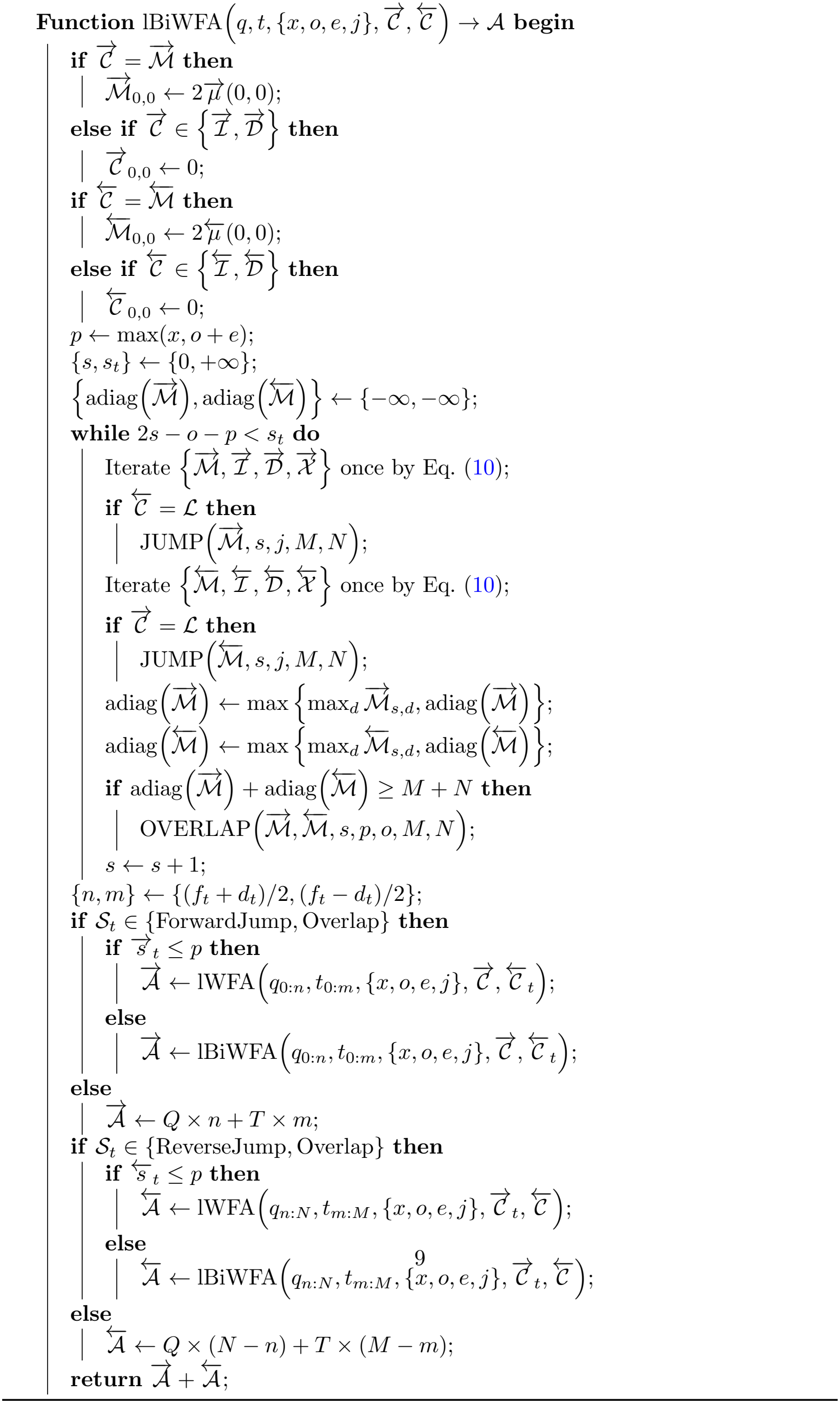

#### Algorithm 3

Jump from the current wavefront.

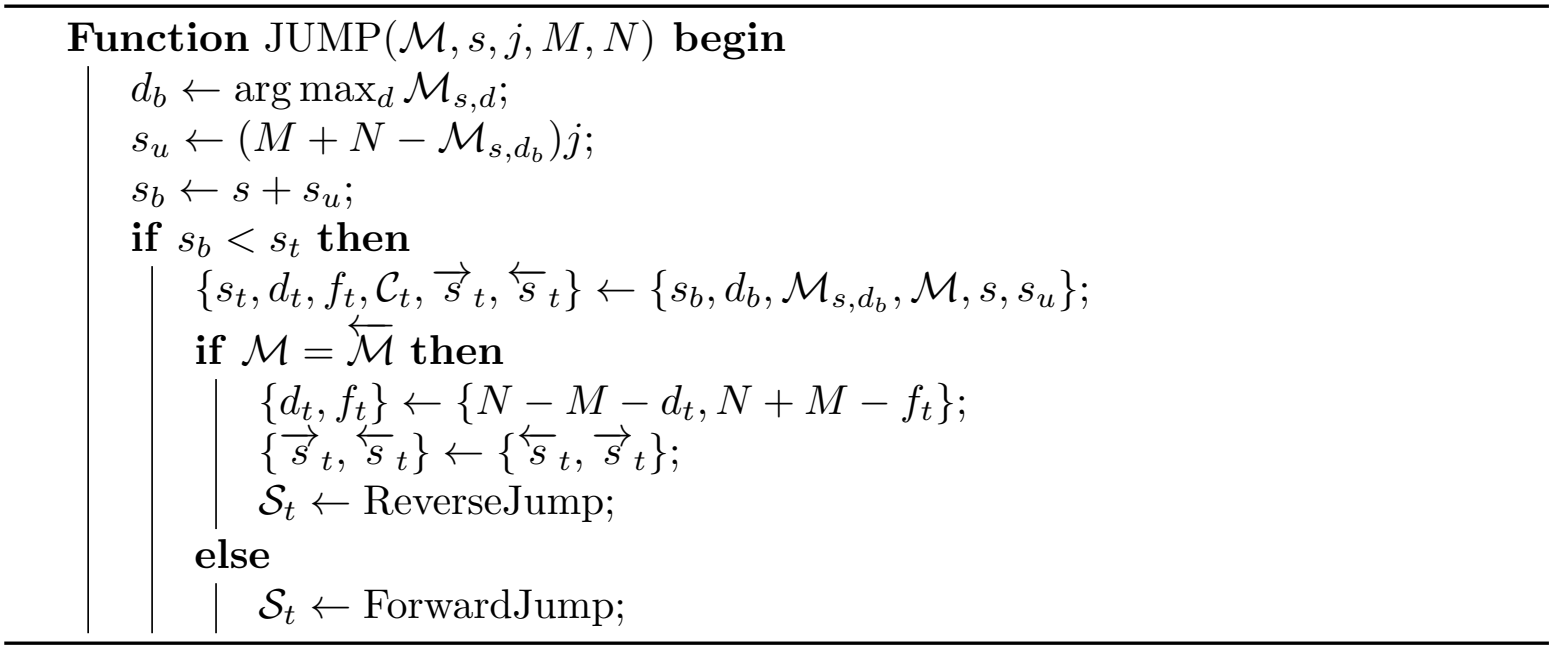

- 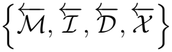 are the wavefront components of the reverse process. To high-light the distinction, denote the wavefront components of the forward process by 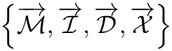.
- Adiag 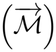 and adiag 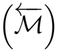 are the most advanced anti-diagonal reached by the forward and reverse wavefront, respectively.
- Different from [21], we set the diagonal 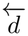 and offset 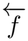 of the reverse process relative to the lower-right corner (*M, N* ) of the DP table. We also define 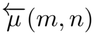 as the length of the longest matching suffix between *t*_0:*M−m*_ and *q*_0:*N−n*_. The original *f, d*, and *µ* (*m, n*) are denoted by 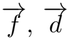 and 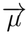(*m, n*) for distinction. The goodness is that the iteration format of the reverse process becomes completely the same as that of the forward one in Eq. (10). The forward and reverse diagonals and offsets are related by

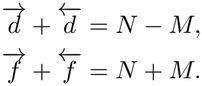
- Denote the unaligned regions in the query and text in the alignment string by *Q* and *T*, respectively.

In Lemmas 1 and 2 in the supplementary material of [21], the authors prove that in the global alignment scope, the optimal score of the partial alignments ending at the same diagonal is non-decreasing with the offsets. Since this result is key to the correctness of the bidirectional wavefront alignment, we extend it to the local alignment scope (refer Appendix A.4 for the proof).

#### Lemma 2

*For* (*m, n*) *∈* [0, *M* ] *×* [0, *N* ], *let* M(*m, n*), I(*m, n*), D(*m, n*) *be the optimal score of the partial local alignment ending at* (*m, n*) *with no tailing restriction, a tailing insertion, and a tailing deletion, respectively. The scoring system is defined in Eq. (6). Then* M(*m, n*), I(*m, n*), *and* D(*m, n*) *are non-decreasing with offsets along each diagonal*.

#### Algorithm 4

Detect overlaps between forward and reverse wavefront and infer the optimal division.

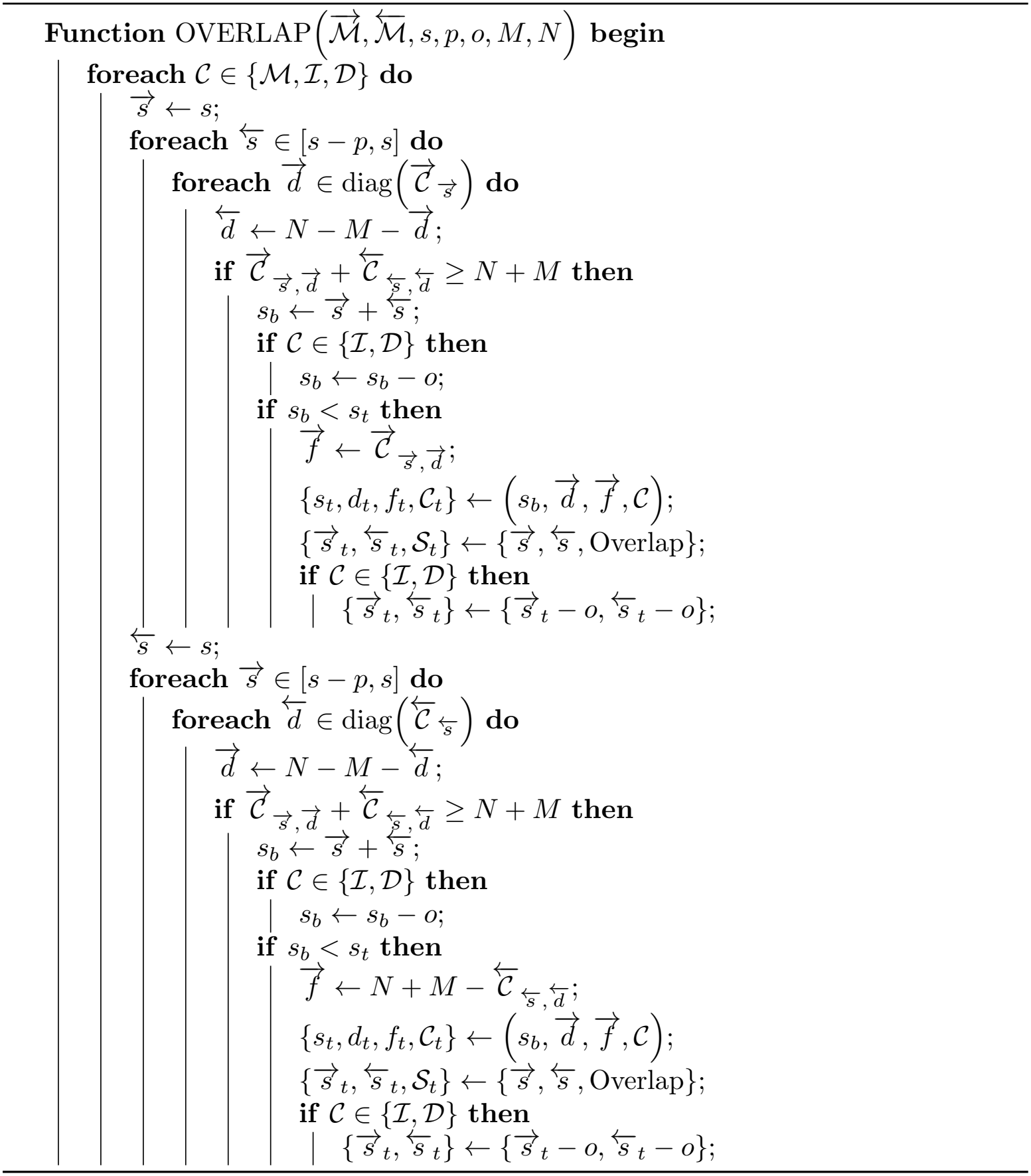

We first prove that Algorithm 2 will not return a score lower than the optimal one (refer Appendix A.5 for the proof).

#### Lemma 3

*In Algorithm 2, s*_*t*_ *≥ s*_*o*_, *where s*_*o*_ *is the optimal alignment score*.

In Lemma 2.1 of [21], the authors show that for the global WFA, the optimal score *s*_*o*_ ≤ *s* if and only if there exist 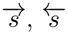 and 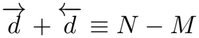 such that 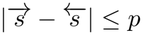 and at least one of the following is true.

- 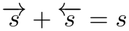 and 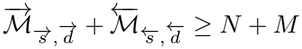 .
- 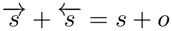 and 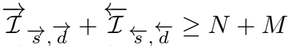 .
- 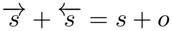 and 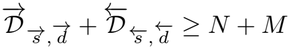.

In the optimal global alignment *A* with score *s*_*o*_, let (*m, n*) on *A* satisfy that the partial alignment from (0, 0) to (*m, n*) along *A* has the score closest to *s*_*o*_*/*2. (*m, n*) may be strictly within (not at boundaries) a deletion, insertion, or otherwise. Those correspond to the three cases in the lemma. In our local alignment scope, there is a fourth case that (*m, n*) is not within the aligned region of the optimal local alignment, but within the unaligned region. This provides a hint to the proof of the correctness of Algorithm 2. We first prove a lemma resembling Lemma 2.1 of [21] (refer Appendix A.6 for the proof).

#### Lemma 4

*Let A be the optimal local alignment with score s*_*o*_. *Let* (*m, n*) *on A satisfy that the partial alignment from* (0, 0) *to* (*m, n*) *along A has the score* 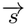 *is closest to s*_*o*_*/*2. *Let* 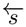 *be the score of the partial alignment from* (*m, n*) *to* (*M, N* ) *along A. If* (*m, n*) *is in the aligned region of A, then* 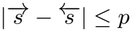.

With the above preparations, we can now prove the correctness of Algorithm 2 (refer Appendix A.7 for the proof).

#### Theorem 1

*Algorithm 2 returns the optimal alignment*.

Resembling Theorem 2.2 in [21], we have Proposition 3 (refer Appendix A.8 for the proof).

#### Proposition 3

*The time and space complexity of Algorithm 2 is O*(*s*_*o*_(*M* + *N* )) *and O*(*s*_*o*_), *respectively, where s*_*o*_ *is the optimal alignment score*.

## 3 Results

### 3.1 Difficulties of local-scope wavefront alignment and our solutions

Extending the wavefront alignment to the local scope presents two difficulties. First, the scoring system in the original WFA paper penalizes errors but does not reward matches [14]. Although this works for global alignment, it cannot be applied directly to local alignment because a zero penalty can always be achieved by aligning nothing. Second, unlike global wavefront alignment, local wavefront alignment has no fixed starting point: any cell in the DP table can serve as a starting point. Tracking wavefront from every starting point would violate the *O*(*sN* ) time and *O*(*s*^2^) space complexity bounds of wavefront alignment. As shown below, these difficulties are closely related: addressing the first also resolves the second.

To address the first difficulty, namely that a vanish alignment incurs no penalty, we assign a linear penalty of *l*_*c*_*J* to the unaligned region, where *l*_*c*_ is the matching score in the classical affine-gap scoring system, and *J* is the length of the unaligned region. To understand this choice, consider aligning two identical sequence of length *L*. In the classical scoring system, a perfect alignment has a score of *l*_*c*_*L*, whereas a vanish alignment has a score of zero. According to this observation, a vanish wavefront alignment should incur a penalty propositional to *l*_*c*_*L* since a perfect one has zero penalty. Accordingly, an unaligned region of length *J* is penalized by *l*_*c*_*J* : the unaligned regions forfeits opportunities to obtain matching rewards. Insertions, deletions and mismatches incur similar opportunity costs. For example, the wavefront penalty *x* for a mismatch equals 2*l*_*c*_ + 2*x*_*c*_, where 2*x*_*c*_ is the mismatch penalty inherited from the classical scoring system, and 2*l*_*c*_ is the opportunity cost of forfeiting a matching reward. A comprehensive discussion of the scoring system is provided in subsection 2.1.

To address the second difficulty, namely the absence of a fixed starting point for local-scope wavefront alignment, we use a dynamic strategy that gradually includes starting points as the score (penalty) increases. For a starting point (*m, n*) in the DP table, the baseline penalty is (*m* + *n*)*l*_*c*_. Because the penalty is non-decreasing, an alignment starting from (*m, n*) cannot have a penalty less than (*m* + *n*)*l*_*c*_. Therefore, alignments starting from (*m, n*) need not be considered when *s <* (*m* + *n*)*l*_*c*_. Fig. 1 illustrates how new starting points are included as needed. Initially, only the single starting point with zero penalty is included. When the score (penalty) *s* reaches *l*_*c*_*J*, starting points with offsets *J* must be included. In general, the offset of new starting points is ⌊*s/l*_*c*_⌋. Although the number of included starting points increases quadratically with *s*, the wavefront bandwidth is bounded above linearly by *s*. In subsection 2.2, we prove that this dynamic strategy yields the optimal local alignment and preserves the time and space complexity of global-scope wavefront alignment.

**Fig. 1:**
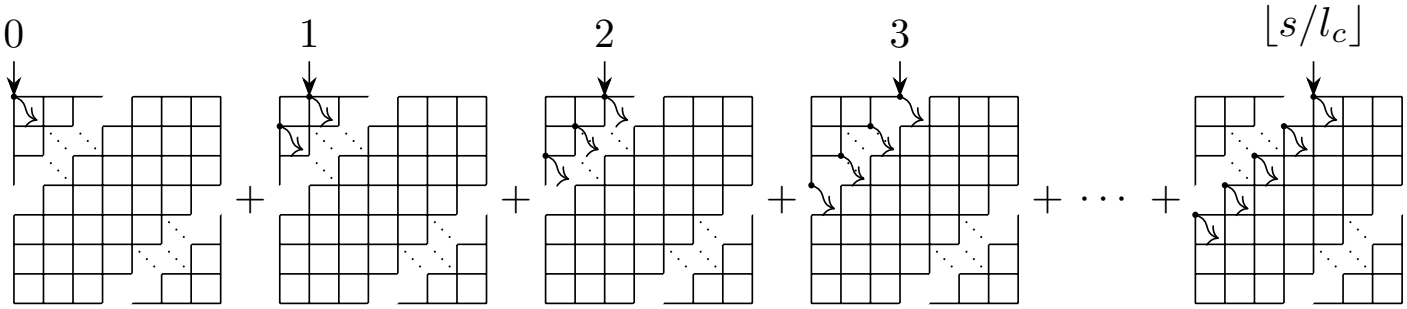
A diagram showing the dynamic process to gradually include starting points for wavefront local scope.

We show that the matching score in the classical scoring system, when applied to the WFA scoring system for the local alignment scope, can be transformed into the linear penalties for unaligned regions. Based on this scoring system, we propose the wavefront alignment for the local scope (lWFA). We prove that lWFA has *O*(*s*^2^) space complexity and *O*(*sN* ) time complexity, which matches the global WFA and significantly surpasses the classical *O*(*N*^2^) space and time complexity of the Smith– Waterman algorithm. We further propose the bidirectional wavefront alignment for the local scope (lBiWFA). lBiWFA improves the space complexity of lWFA from *O*(*s*^2^) to *O*(*s*) while maintaining the time complexity as *O*(*sN* ). Benchmarks on simulated data demonstrate that lWFA and lBiWFA outperform the exact local alignment libraries such as Parasail and SeqAn in both time and space efficiency when aligning long reads with moderate error rates.

### 3.2 Benchmarks

Although there exist many highly efficient pairwise-alignment libraries, such as Parasail [8], SeqAn [24], block-aligner [25], Edlib [26], BitPAl [13], wfalm [20], WFA2-lib [14, 21], Gaba [27], and KSW2 [28, 29], most of them are either heuristic or do not support the local alignment scope with affine gap penalties (Fig. 2). Only Parasail and SeqAn have exact methods that support the local alignment scope with affine gap penalties. We benchmark lWFA and lBiWFA against the diagonal method by Wozniak (ParaDiag) [30], the striped approach by Farrar [11], the prefix-scan algorithm (ParaScan) [12], and the SeqAn library [24].

**Fig. 2:**
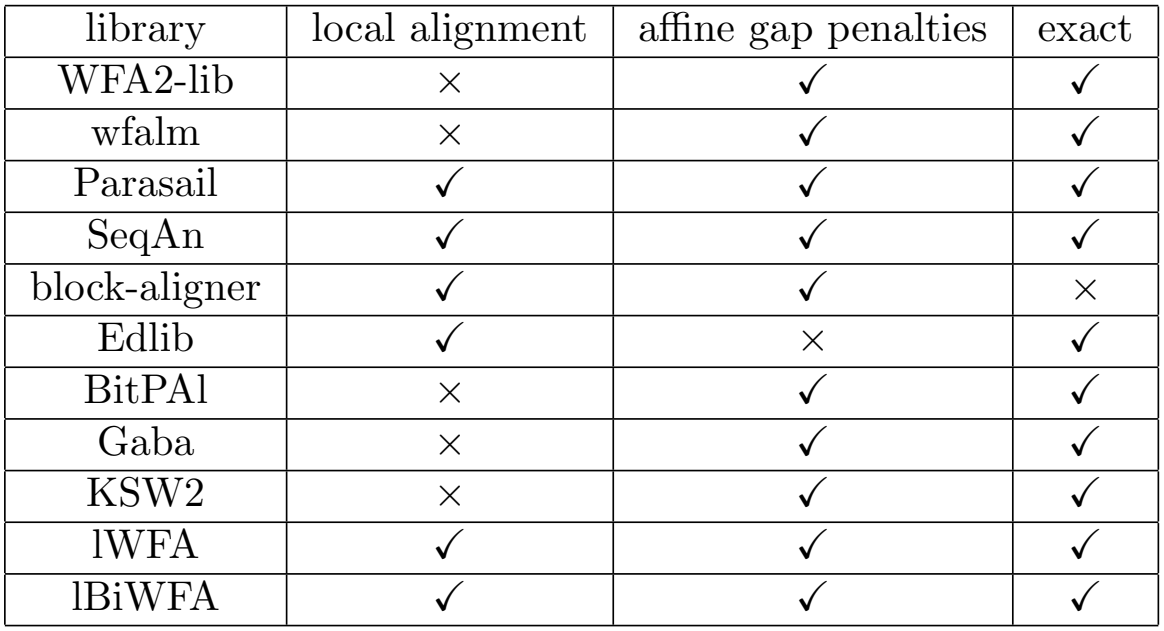
The feature grid of the pairwise-alignment libraries.

We benchmark on both simulated and real data on the Bowtie2 scoring system.

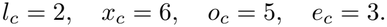

We first test on simulated data. We generate 100 random DNA reads for each combination of length *l* in 100bp, 1kbp, 10kbp and error rate *r* in 1%, 5%, 10%. To make the simulated reads suitable for the local alignment scope, we mutate *l* × *r* bases at both ends of the reads. The peak space usage of 100bp and 1kbp reads is negligible compared to the benchmark overhead for all tested libraries (Figs. 3A and 3B). For 10kbp reads, the peak space usage of lBiWFA is the lowest over all error rates, and is negligible compared to the benchmark overhead (Fig. 3C). The peak space usage of lWFA scales well with the read length but not the error rate. SeqAn and Parasail use more memory as the read length increasing. Their peak space usage are irrelevant to the error rate.

**Fig. 3:**
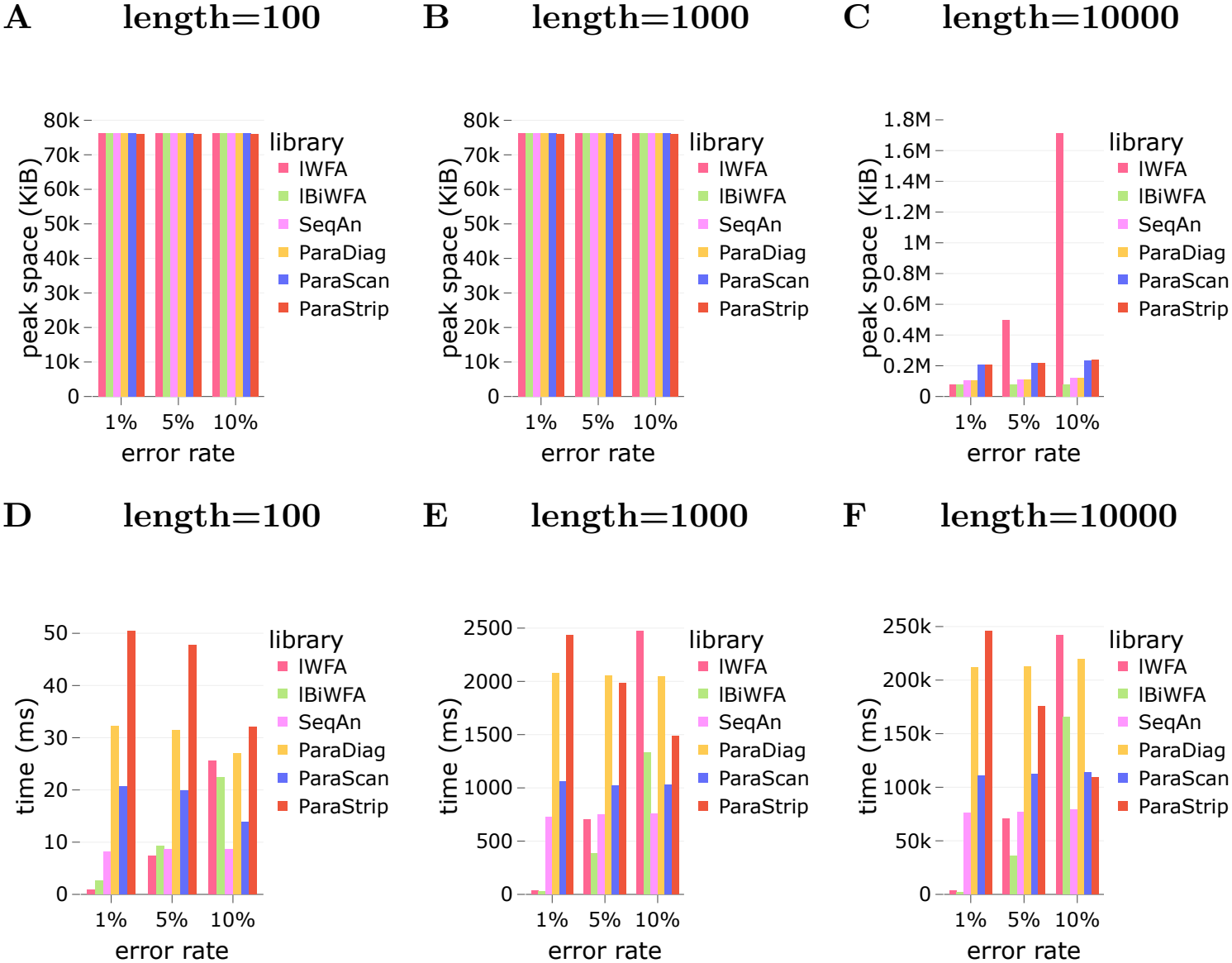
Peak space **(A,C,E)** and time **(B,D,F)** usage of lWFA, lBiWFA, SeqAn, ParaDiag, ParaStrip, ParaScan on aligning 100 simulated sequences for each combination of length in 100bp, 1kbp, 10kbp and error rate in 1%, 5%, 10%.

lWFA and lBiWFA are faster than SeqAn and Parasail for 100bp reads of 1% error rate, as well as 1kbp and 10kbp reads of 1% and 5% error rates. lWFA, lBiWFA and SeqAn are faster than Parasail for 100bp reads of 5% error rate. For reads of 10% error rate, lWFA and lBiWFA are slower than SeqAn and Parasail.

We then select 100,000 Illumina HiSeq sequences of 100bp (Accession number ERR091574) [31]. The peak space usage is negligible compared with the benchmark overhead for all tested libraries (Fig. 4A). lWFA is the fastest, and lWFA and lBiWFA are much faster than SeqAn and Parasail (Fig. 4B).

**Fig. 4:**
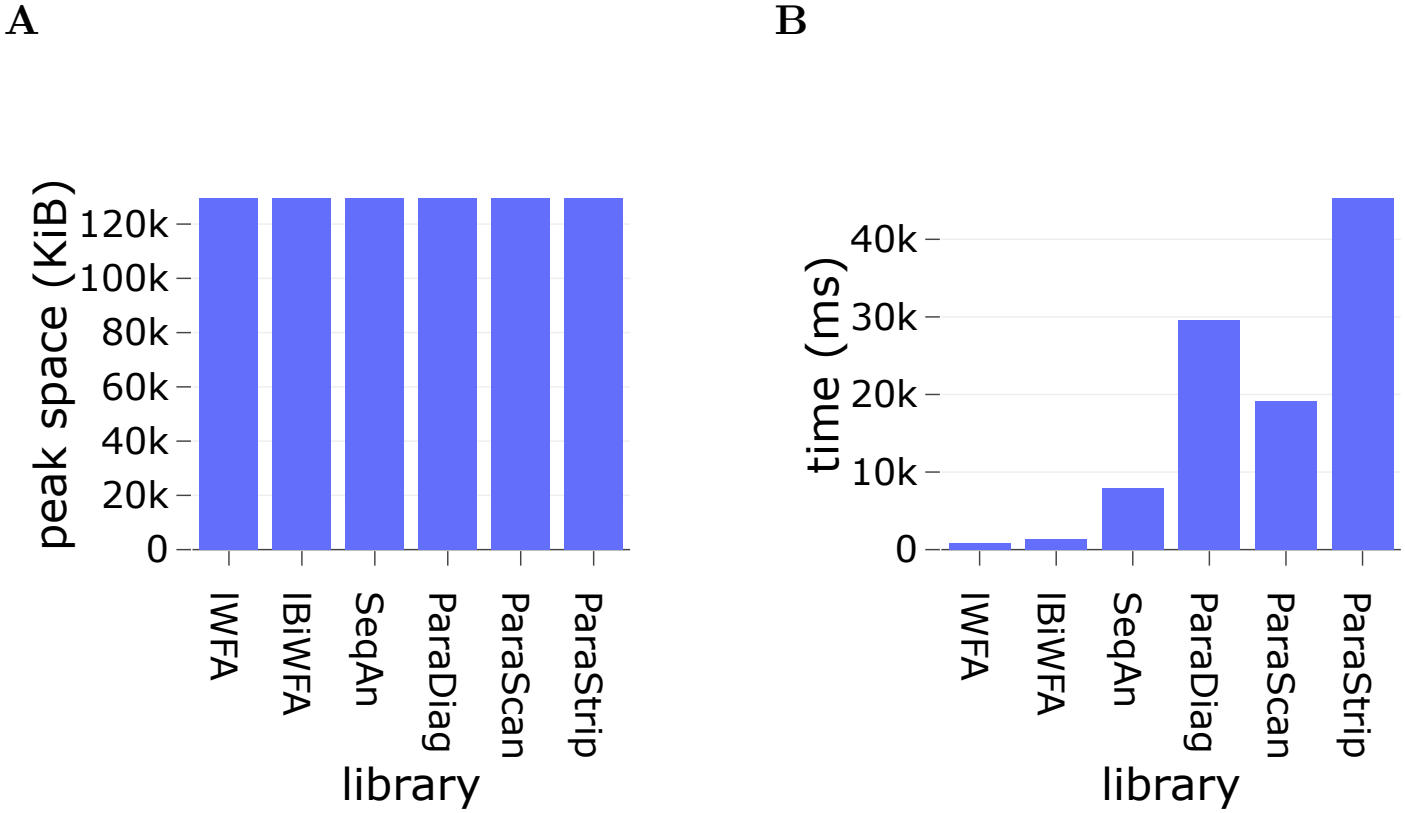
Peak space **(A)** and time **(B)** usage of lWFA, lBiWFA, SeqAn, ParaDiag, ParaStrip, ParaScan on aligning 100,000 Illumina HiSeq sequences of 100bp.

We finally select 1,000 ONT MinION sequences of 10 ∼ 20kbp (Accession number ERR3278877) [31]. We filter out reads with mapping qualities lower than 30 or mapping coverages lower than 50% (possibly chimeric reads). lBiWFA has the least peak space usage (Fig. 5A). However, lWFA uses much more memory than SeqAn and Parasail even if its *O*(*s*^2^) space complexity is lower than *O*(*N*^2^) of SeqAn and Parasail. We conjecture that this is because our current implementation is naive and lacks production-grade memory optimization strategies. lBiWFA is the fastest, followed by lWFA. SeqAn is slightly slower than lWFA (Fig. 5B).

**Fig. 5:**
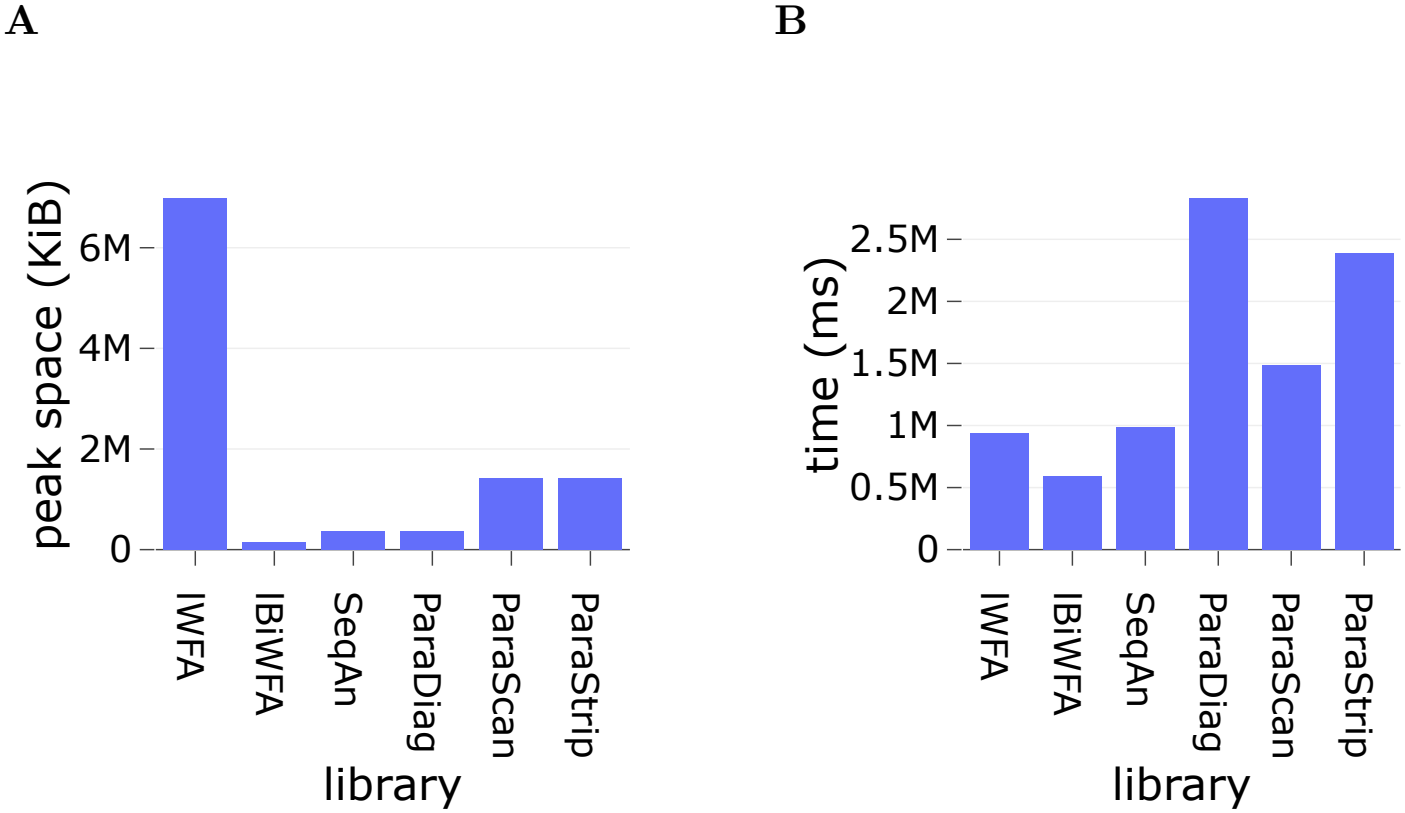
Peak space **(A)** and time **(B)** usage of lWFA, lBiWFA, SeqAn, ParaDiag, ParaStrip, ParaScan on aligning 1,000 ONT MinION sequences of 10∼20kbp.

In summary, for reads with *<* 1% error rate (say NGS reads with 0.1% ∼ 1% error rate), lWFA and lBiWFA are greatly faster than SeqAn and Parasail. For reads with *<* 5% error rate (say ONT reads with 3% ∼ 5% error rate), lBiWFA has an obvious advantage. However, for reads with *>* 10% error rate (say reads from legacy long-read sequencing platforms), SeqAn and Parasail are faster than lWFA and lBiWFA.

According to the time benchmarks in Fig. 5B (ONT MinION reads) and Figs. 3D-3F (simulated reads), lBiWFA has a higher tolerance to error rates than lWFA. To clarify the reason, we use both lWFA and lBiWFA to align the same 100bp read with 5% error rate and 10bp unaligned regions at both ends. The DP cells visited by lWFA and lBiWFA are shown in Fig. 6. Compared with lWFA, lBiWFA requires only half the score at each direction to meet in the middle. This greatly reduces the number of DP cells visited by lBiWFA, thereby reducing the effective wavefront bandwidth. The jumping steps of lWFA and lBiWFA lead to dense triangles of visited cells in the start and end corners of the DP table. These are additional expenses of the local-scope wavefront alignment compared with the global-scope one.

**Fig. 6:**
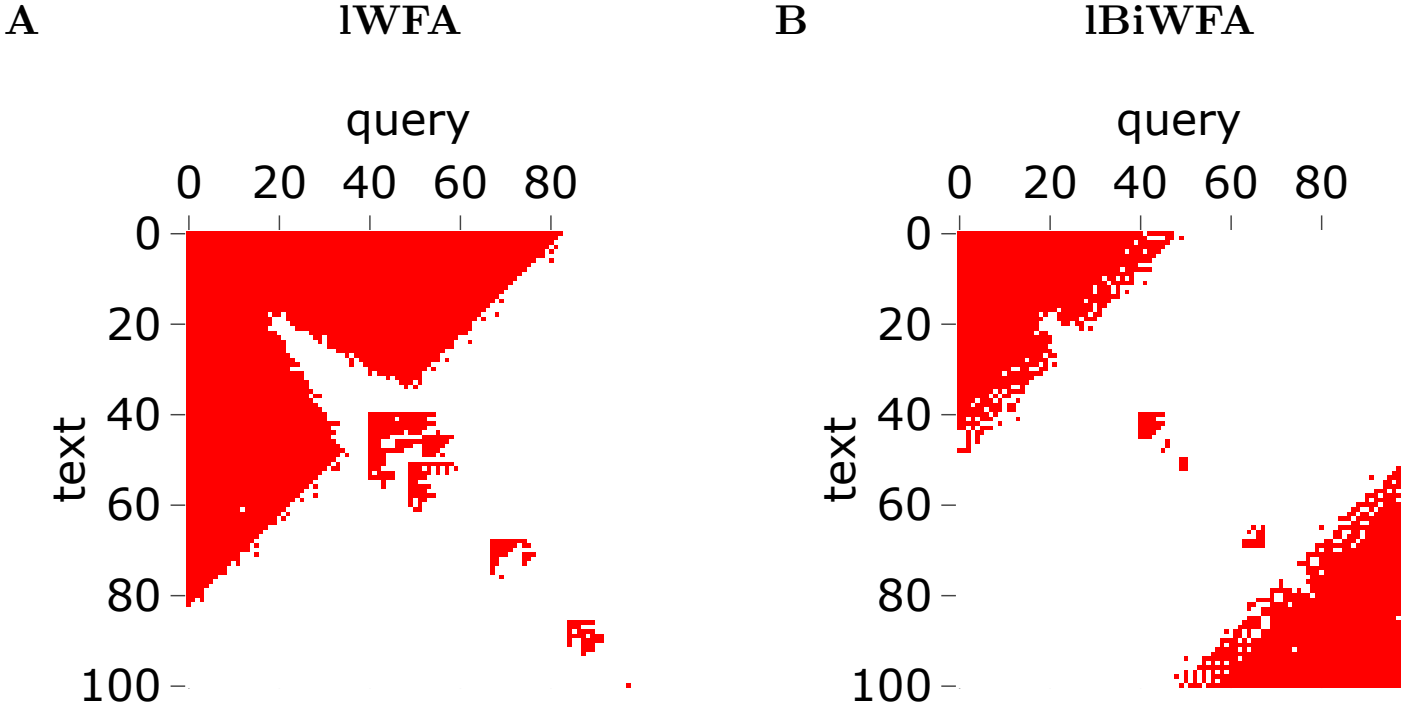
Visited DP cells of lWFA **(A)** and lBiWFA **(B)** for aligning the same 100bp read with 5% error rate and 10bp unaligned regions at both ends.

## 4 Discussion

The semi-global implementation in WFA2-lib does not penalize the unaligned region [14, 21]. Instead, it presets an upper bound on the size of the unaligned region, which is heuristic in nature. In contrast, our free-end corrected scoring system not only is equivalent to the classical one, but also enables lWFA and lBiWFA to progressively extend the unaligned region and obtain the exact optimal local alignment. More-over, our progressive-extension method also works in the semi-global scope by simply restricting the unaligned region in the text or the query but not both. Our method does not preset an upper bound on the size of the unaligned region, thereby being exact instead of heuristic.

lWFA and lBiWFA achieve a favorable theoretical time complexity of *O*(*sN* ). However, they face several practical limitations. For high-error reads, Parasail and SeqAn outperform lWFA and lBiWFA due to the large score penalty *s* in the theoretical time complexity *O*(*sN* ) of lWFA and lBiWFA. When the error count or the unaligned read region scales proportionally with the read length *N*, the *O*(*sN* ) time complexity degenerates to *O*(*N*^2^). Moreover, lWFA and lBiWFA must manage the sparse structure of the wavefront, incurring additional overhead. Our benchmarks use the Bowtie2 scoring system. However, the speed of lWFA and lBiWFA is sensitive to the actual scoring system used. This is because in Algorithms 1 and 2, the diagonal range of the wavefront expands whenever 0 = *s* (mod *j*). Since the jumping penalty *j* in the lWFA scoring system corresponds to the match bonus *l*_*c*_ in the classical one, the wave-front bandwidth is roughly proportional to the ratio of the total error penalty to the single match bonus. For the global-scope wavefront alignment, its bandwidth is lower-bounded by *s/e* with *e* = 2*e*_*c*_ + *l*_*c*_ being the gap extension penalty (see Eq. (6)). In practice, *e* ≫ *j*, which makes the wavefront alignment much faster in the global scope than the local one.

Our current implementations of lWFA and lBiWFA are far from perfect. We store DNA sequences as plain strings, which consumes 8 bits per base. Since there are only 4 types of bases, it would be more space-efficient to store each base using 2 bits. Another benefit of the 2-bit representation is that we can pack up to 32 bases into a single 64-bit machine word. In this way, 32 bases can be compared in parallel during the perfect matching extension. Our implementation always use 32-bit integer and does not adapt the bit width to the read length. For short reads, using 16-bit integer is more efficient.

## 5 Conclusions

The current scoring system of wavefront alignment only penalizes errors and does not reward matchings. This prevents its application in the local alignment scope because one may always achieve zero penalty by aligning nothing in the local alignment scope. We extend the scoring system of wavefront alignment by introducing penalties for the unaligned regions in the local alignment scope. We prove that this new scoring system is equivalent to the classical affine-gap scoring system of the local alignment scope used by the Smith–Waterman algorithm.

We design the local wavefront alignment algorithm (lWFA) by including a jumping step to skip unaligned regions at both read ends. lWFA preserves the *O*(*sN* ) time and *O*(*s*^2^) space complexity of its global version. By applying the divide-and-conquer design to lWFA, we obtain the bidirectional version of lWFA (lBiWFA). lBiWFA preserves the *O*(*sN* ) time complexity of lWFA and has a *O*(*s*) space complexity. We provide strict proofs for the correctness and complexity of lWFA and lBiWFA.

In the global scope, the bidirectional version of wavefront alignment is slower because of the overhead in its recursive calling. However, in the local scope, lBiWFA is generally faster than lWFA (except for short reads with very low error rates), and has a higher tolerance to error rates. This is because jumping step in the local alignment scope introduces a dense triangle of visited cells at corners of the DP table (Fig. 6). lBiWFA distributes the penalties to both directions, which divides the single large triangle of lWFA into two smaller ones. This explains why lBiWFA is faster than lWFA for reads with high error rates.

Our benchmarks suggest to use lBiWFA over lWFA because our current implementation of lWFA has a large memory footprint, and lWFA only outperforms lBiWFA for short reads with very low error rates. lBiWFA has a negligible memory footprint for aligning reads up to 20kbp. It is faster than the current local alignment libraries such as SeqAn and Parasail for aligning NGS short reads with 0.1% ∼ 1% error rate and ONT long reads with 3% ∼ 5% error rate. For reads sequenced by legacy long-read platforms with 10% ∼ 15% error rate, SeqAn and Parasail is faster but lBiWFA still has a competitive speed. If the memory usage is the major consideration, one may choose lBiWFA over SeqAn and Parasail.

## Declarations

### Funding

This research was supported by the National Natural Science Foundation of China (32330016 to QW), the National Key R&D Program of China (2022YFC3400200 to QW) and the Shanghai Sailing Program (21Z510202245 to JL). The funders had no role in study design, data collection and analysis, decision to publish, or preparation of the manuscript.

### Consent for publication

Not applicable.

### Ethics approval and consent to participate

Not applicable.

### Competing interests

The authors declare no competing interests.

### Availability of data and materials

The source code of the lWFA and lBiWFA is publicly available under the MIT license at the GitHub repository: https://github.com/ljw20180420/wavefront_local. A PyPI package named wavefront local is available for the Python bindings of our algorithms.

### Authors’ contributions

J.L. designs the algorithm. J.L. writes the package. H.Z. tests the package. J.L., H.Z., L.J. and Q.W. write the paper.

## Acknowledgements

Not applicable.

## Appendix A Proofs

### A.1 The proof of Lemma 1

*Proof* We first prove for *ℐ*_*s,d*_. Assume by induction that all wavefront components so far satisfy the lemma assumption. Prove by contradiction that there exists a partial alignment ending at diagonal *d* with score *s* and a tailing insertion but has a larger offset 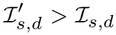. By Eq. (10), 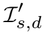 has two possible sources.

- 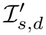 comes from an insertion with offset 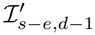. By Eq. (10), we have ℐ_*s,d*_ ≥ ℐ_*s−e,d−*1_ + 1. Since 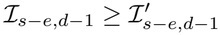 by induction, we have 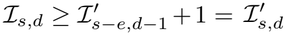, conflicts.
- 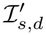 comes from a non-insertion with offset 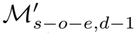. By Eq. (10), we have ℐ_*s,d*_ ≥ ℳ_*s−o−e,d−*1_ + 1. Since 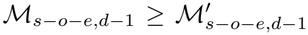 by induction, we have 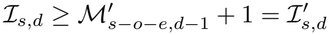, conflicts.

The proof for *D*_*s,d*_ is similar to that for *ℐ*_*s,d*_.

We next prove for *ℳ*_*s,d*_. Assume by induction that all wavefront components so far satisfy the lemma assumption. Prove by contradiction that there exists a partial alignment ending at diagonal *d* with score *s* but has a larger offset 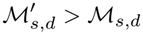. Assume that *ℳ*_*s,d*_ and 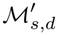 come from *X*_*s,d*_ and 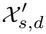 by the perfect matching extension along diagonal *d*, respectively. By Eq. (10), 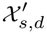 has four possible sources.

- 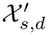 comes from a jumping of size *s/j*. This implies 0 = *s* (mod *j*). By Eq. (10), 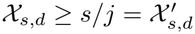, which implies 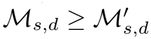, conflicts.
- 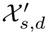 comes from an insertion with offset 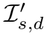. By Eq. (10), we have *X*_*s,d*_ ≥ ℐ_*s,d*_. Since 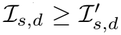 by induction, we have 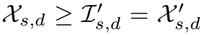, which implies 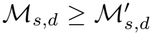, conflicts.
- 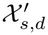 comes from a deletion. The conflict is similar to that of an insertion.
- 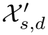 comes from a mismatch with offset 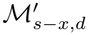 By Eq. (10), we have *X*_*s,d*_ ≥ ℳ_*s−x,d*_ + 2. Since 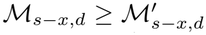 by induction, we have 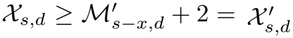, which implies 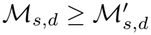, conflicts. □

### A.2 The proof of Proposition 1

*Proof* We first prove the case for 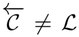. Algorithm 1 returns when 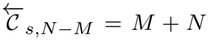 . By Lemma 1, this happens if and only if there exists an alignment ending at (*M, N* ) with a tailing 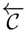. Since *s* is the minimal score such that 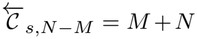, the alignment *A* traced from 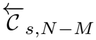 must be optimal.

We next prove the case for 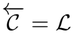. Algorithm 1 exhausts the alignments with the tailing jumping from *ℳ*_*s,d*_, which are wavefront components by Lemma 1. The proof is then straight forward by noting that compared with other partial alignments with the same score, those corresponding the wavefront components have the maximal offsets, thereby minimal penalty at the tailing jumping. □

### A.3 The proof of Proposition 2

*Proof* The diagonal range increases at most by 2 for each *s* due to the iterations of *ℐ*_*s,d*_ and *D*_*s,d*_, and is bounded by [*−s, s*]. When 0 = *s* (mod *j*), a jumping of size *f* = *s/j* has a penalty exactly equal to *s*, and Algorithm 1 updates all diagonals ranging from max(*−f, f −* 2*M* ) to min(*f*, 2*N − f* ) by 2 with a least offset *f* . The diagonals updated by the jumping are bounded by [*−f, f* ] *⊂* [*−s, s*]. Thus, each iteration of *s* has a space complexity *O*(*s*). Algorithm 1 iterates *s* until *s*_*t*_ *≤ s*. By Proposition 1, *s*_*t*_ = *s*_*o*_ at the end of the score iteration. Thus, there are totally *s*_*o*_ iterations, thereby the space complexity is 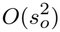. Note that *X*_*s,d*_ *≥ ℳ*_*s−x,d*_ +2. Therefore, the perfect matching extensions for *s* = *y, x* + *y*, 2*x* + *y, · · ·* with 0 *≤ y < x* do not overlap along any diagonal. This implies that the total extensions are bounded up to a scaling factor by *xs*_*o*_(*M* + *N* ). Therefore, the time complexity of Algorithm 1 is *O*(*s*_*o*_(*M* + *N* )). □

### A.2 The proof of Lemma 2

*Proof* We only prove the case for M since the proofs of I and D are similar. For each (*m, n*) *∈* [0, *M* ] *×* [0, *N* ], define the score function S_*m,n*_(*·, ·*) on [0, *M* ] *×* [0, *N* ] as follows.

- If *m*^*′*^ *< m* or *n*^*′*^ *< n*, then S_*m,n*_(*m*^*′*^, *n*^*′*^) = (*m*^*′*^ + *n*^*′*^)*j*.
- Otherwise, S_*m,n*_(*m*^*′*^, *n*^*′*^) is the optimal score of the global alignment from (*m, n*) to (*m*^*′*^, *n*^*′*^), plus (*m* + *n*)*j*.

By Lemma 1 in the supplementary material of [21], S_*m,n*_ is non-decreasing with offsets along each diagonal in the region [*m, M* ] *×* [*n, N* ]. The non-decreasing property can be extended to the whole region [0, *M* ] *×* [0, *N* ] by noting that *e* = 2*e*_*c*_ + *l*_*c*_ *≥ l*_*c*_ = *j* in Eq. (6). By definition,

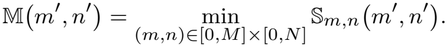

Since each S_*m,n*_ is non-decreasing along diagonals, so is M. □

### A.5 The proof of Lemma 3

*Proof* Initially, *s*_*t*_ = +*∞ > s*_*o*_. Algorithm 2 updates *s*_*t*_ by JUMP and OVERLAP. We first discuss JUMP. By definition, there exists a partial alignment ending at diagonal *d*_*b*_ with offset 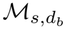 and score *s*. JUMP completes it by a tailing jumping penalized by 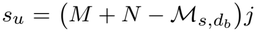, leading to a complete local alignment with score *s*_*b*_ = *s* + *s*_*u*_. By definition, *s*_*b*_ *≥ s*_*o*_. Since JUMP updates *s*_*t*_ by *s*_*b*_, we have *s*_*t*_ *≥ s*_*o*_.

We next discuss OVERLAP. We only prove the case with 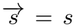 since the case with 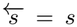 is similar. Let 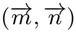 and 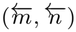 be the DP cells corresponding to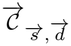 and 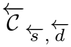, respectively. By the definition of 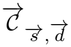, there exists a forward partial alignment 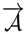 ending at 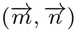 with score 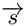 and a tailing *C*. Similarly, for 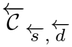, there exists a reverse partial alignment 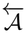 ending at 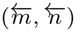 with score 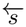 and a tailing *C*. By Lemma 2, the score 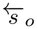 of the optimal reverse partial alignment 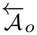 ending at 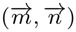 with a tailing *C* satisfied 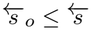. Joining 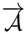 and 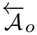, there is

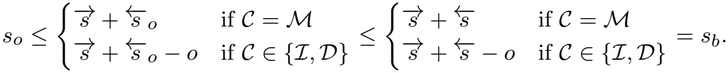

Since OVERLAP updates *s*_*t*_ by *s*_*b*_, we have *s*_*t*_ *≥ s*_*o*_. □

### A.6 The proof of Lemma 4

*Proof* There are two cases.

- If (*m, n*) is not strictly within a deletion or insertion, then 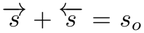. Since 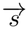 is closest to *s*_*o*_*/*2 and the score increment of each step does not exceed *p*, there is 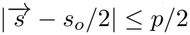. Thus,

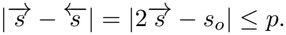
- If (*m, n*) is strictly within a deletion or insertion, then 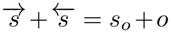. If 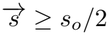, then

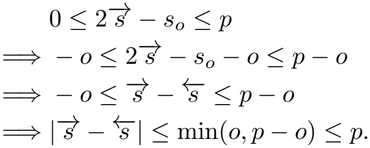

Otherwise, 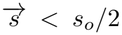. Assume without loss of generality that (*m, n*) is strictly within an insertion. Then the score increment from (*m, n*) to (*m, n* + 1) is *e* because (*m, n* − 1) to (*m, n*) is also an insertion. Since the score 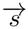 at (*m, n*) is closer to *s*_*o*_*/*2 than that at (*m, n* + 1), we have 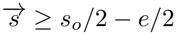. Thus,

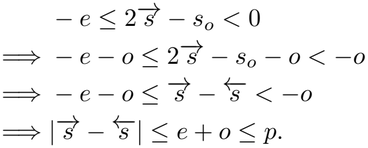

### A.7 The proof of Theorem 1

*Proof* We first discuss the case for 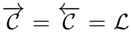. Divide the optimal local alignment *A* into three regions.

- (0, 0) to (*m*^*′*^, *n*^*′*^) is the heading jumping with score 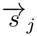.
- (*m*^*′*^, *n*^*′*^) to (*m*^*′′*^, *n*^*′′*^) is the aligned region with score *s*_*a*_.
- (*m*^*′′*^, *n*^*′′*^) to (*M, N* ) is the tailing jumping with score 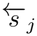.

Then the optimal score *s*_*o*_ satisfies 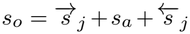 . A call of lBiWFA leads to three cases.

- If 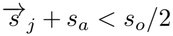, then for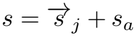, the while loop condition is guaranteed by Lemma 3 as follows.

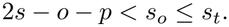

After iterating 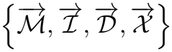 once by Eq. (10) for *s*, there 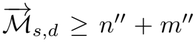 by definition since *A* reaches (*m*^*′′*^, *n*^*′′*^) with score *s*. Let (*m, n*) correspond to 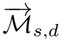. Define 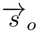 as the optimal score of the alignment between *t*_0:*m*_ and *q*_0:*n*_. Then 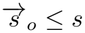 by the definition of 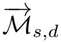. Thus,

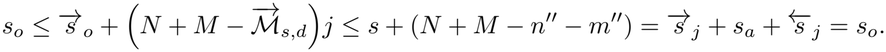

This implies that

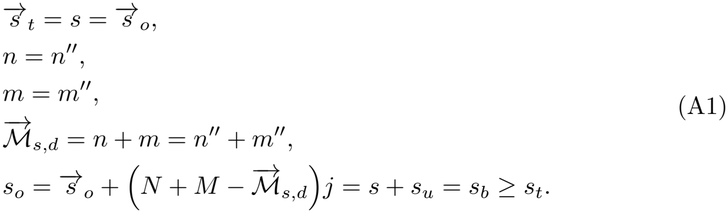

Lemma 3 together with Eq. (A1) implies *s*_*t*_ = *s*_*o*_. If Algorithm 2 can get the optimal alignment between 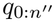 and 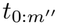 by the recursive call of lBiWFA (or lWFA based on whether 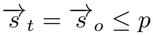 or not), then since 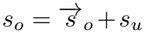, it will return an optimal alignment between *q* and *t*.

- If 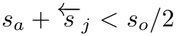, then the discussion is similar to that of 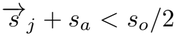 but in the reverse direction.
- If neither 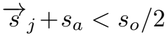 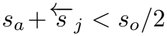, then both 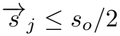 and 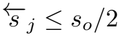.

Therefore, (*m, n*) on *A* defined in Lemma 4 is within the aligned region, *i*.*e*. between (*m*^*′*^, *n*^*′*^) and (*m*^*′′*^, *n*^*′′*^), and 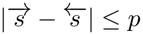. Let *C* ∈ { ℳ, ℐ, *D*} indicate whether (*m, n*) is strictly within a deletion, insertion or not. Without loss of generality, assume 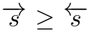 . As long as *s*_*t*_ *> s*_*o*_, the while loop condition for 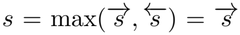 is satisfied by

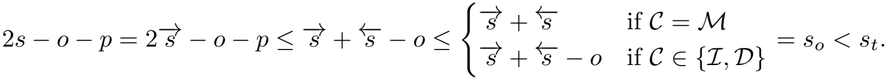

After iterating 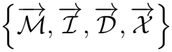 and 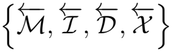 once by Eq. (10), we have by the definition of wavefront that

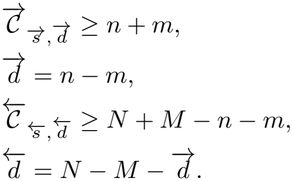

In OVERLAP, the condition 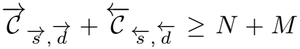 is satisfied. Let (*m*_0_, *n*_0_) correspond to 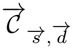 . By the definition of 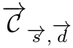, the optimal score 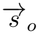 of aligning 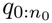 and 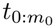 with a tailing *C* satisfies

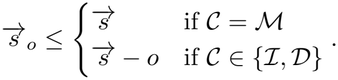

Let 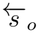 be the optimal score of aligning 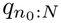 and 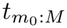 with a heading C. By Lemma 2 and the definition of 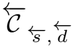,

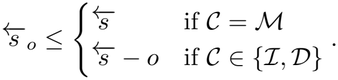

Thus,

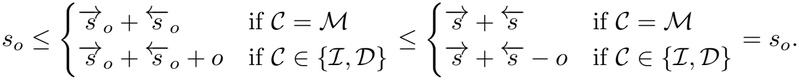

This implies that

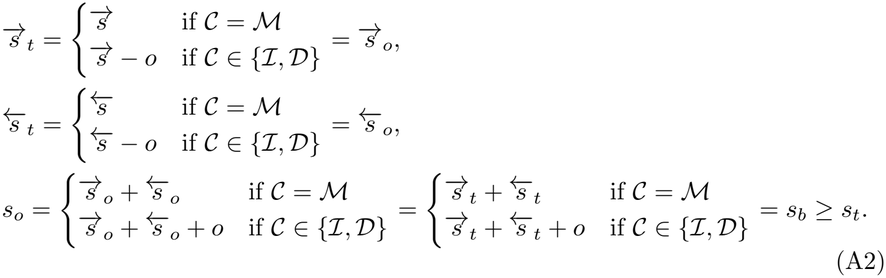

Lemma 3 together with Eq. (A2) implies *s*_*t*_ = *s*_*o*_. Therefore, if Algorithm 2 can get the optimal alignments of 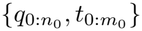 and 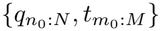 by the recursive calls of lBiWFA (or lWFA based on 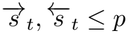 or not), then since

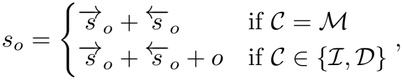

it will return an optimal alignment between *q* and *t*.

The cases for 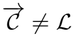 and/or 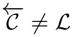 are similar.

We then show that the recursive of lBiWFA has a finite depth. Note that JUMP removes the unaligned region from the alignment, which is called at most twice (one for each end). For OVERLAP, we have

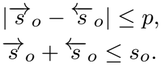

As long as *s*_*o*_ *> p*, there is

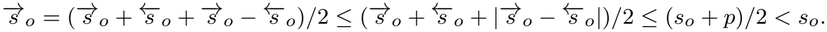

Similarly, 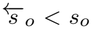 . Therefore, lBiWFA will progressively divide the alignment into smaller ones until *s*_*o*_ *≤ p*. Thus,

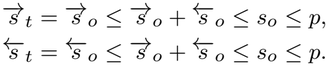

Consequently, lWFA will be called instead of lBiWFA, which returns immediately without further recursions.

Finally, because lWFA returns the optimal alignment, lBiWFA must also return the optimal alignment by induction. □

### A.8 The proof of Proposition 3

*Proof* The space complexity is *O*(*s*_*o*_) because by the proof of Proposition 2, the bandwidth of lWFA is restricted to [*−s, s*], and *s* is restricted by the loop condition 2*s − o − p < s*_*t*_ (when the loop exits, *s*_*t*_ = *s*_*o*_ by Theorem 1).

Now we prove the time complexity. Let *l* = *M* + *N* and *T* (*l, s*_*o*_) be the maximal execution time of lBiWFA and lWFA. For *s*_*o*_ *≤* 3*p*, there exists a constant *c*_2_ such that *T* (*l, s*_*o*_) *≤ c*_2_*l*. This is because the total recursion depth is bounded due to *s*_*o*_ *≤* 3*p*, and in each recursive call of lBiWFA, the lWFA iterations from both directions have the time complexity *O*(*s*_*o*_*l*) by Proposition 2, thereby bounded by *c*_1_*s*_*o*_*l* for large enough constant *c*_1_. A call of lBiWFA leads to three cases.

- In the first case, lBiWFA divides the alignment at (*m, n*) by OVERLAP. Let *C* ∈ { ℳ, ℐ, *D*} indicate whether (*m, n*) is strictly within a deletion, insertion or not. In this case, lBiWFA recursively calls lBiWFA (or lWFA) twice with execution time 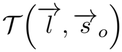 and 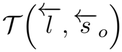, respectively, where

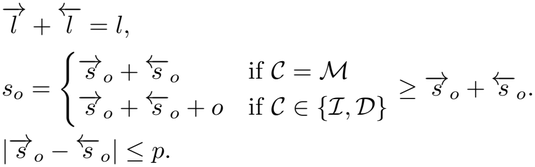

Because the lWFA iterations from both directions are bounded by *c*_1_*s*_*o*_*l*, we have

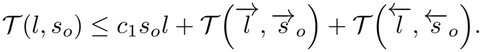

Prove by induction that *T* (*l, s*_*o*_) ≤ 3*c*_1_*s*_*o*_*l* + *c*_2_*l*. Firstly, it is obvious for *s*_*o*_ ≤ 3*p* by *T* (*l, s*_*o*_) ≤ *c*_2_*l*. For *s*_*o*_ *>* 3*p*, because 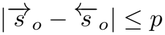 and 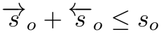, we have

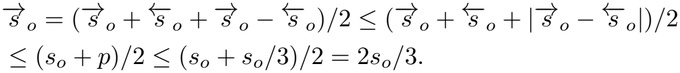

Similarly, 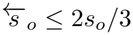. Thus, by induction,

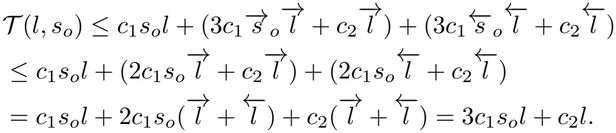

- In the second case, lBiWFA removes the tailing jumping by JUMP. In this case, lBi-WFA recursively calls lBiWFA (or lWFA) only once with execution time 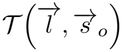 and process a tailing jumping of size 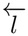. By the first case in the proof of Theorem 1, 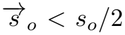. Without loss of generality, we also assume the processing time of the tailing jumping of size 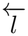 is bounded by 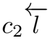 as well. Thus, by induction,

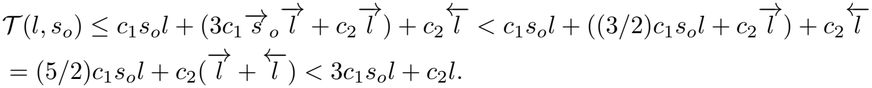
- In the third case, lBiWFA removes the heading jumping by JUMP, which is similar to the second case but from the reverse direction.

## Notes

### Competing Interest Statement

The authors have declared no competing interest.

